# Early retinal microglial activation and ganglion cell dysfunction following severe traumatic brain injury in mice

**DOI:** 10.64898/2026.06.26.734783

**Authors:** Loretta Péntek, Endre Czeiter, Krisztina Amrein, András Szentiványi, Balázs Kovács, Boglárka Balogh, Gergely Szarka, Béla Völgyi, Tamás Kovács-Öller

## Abstract

Traumatic brain injury (TBI) induces rapid neuroinflammatory responses not only in the brain but also in anatomically and immunologically connected central nervous system (CNS) compartments, including the retina. In our study, we investigated retinal microglial activation, retinal ganglion cell (RGC) calcium dynamics, and caspase-3 activation in adult mice subjected to severe traumatic brain injury using the Marmarou impact-acceleration model at 24 and 48 h post-injury. Carrying out Ca²⁺-imaging, immunohistochemistry, and ex vivo time-lapse microscopy, we found robust microglial activation in both the superficial and deep retinal layers following TBI, accompanied by increased microglial motility. RGCs exhibited a transient surge in degeneration-induced spontaneous activity at 24 h, followed by a marked reduction below control levels at 48 h, consistent with early degenerative changes. Activated caspase-3 levels were significantly elevated in both microglia and other retinal cell types at both time points, indicating ongoing apoptotic effects. Together, these findings demonstrate that TBI rapidly triggers inflammatory and apoptotic mechanisms in the retina, which are detectable within the first 48 hours. Our results highlight the retina as a sensitive indicator of early CNS pathology after traumatic injury and underscore the potential of retinal analysis for monitoring TBI-induced neurodegeneration for future clinical implementation.

**Graphical abstract:** 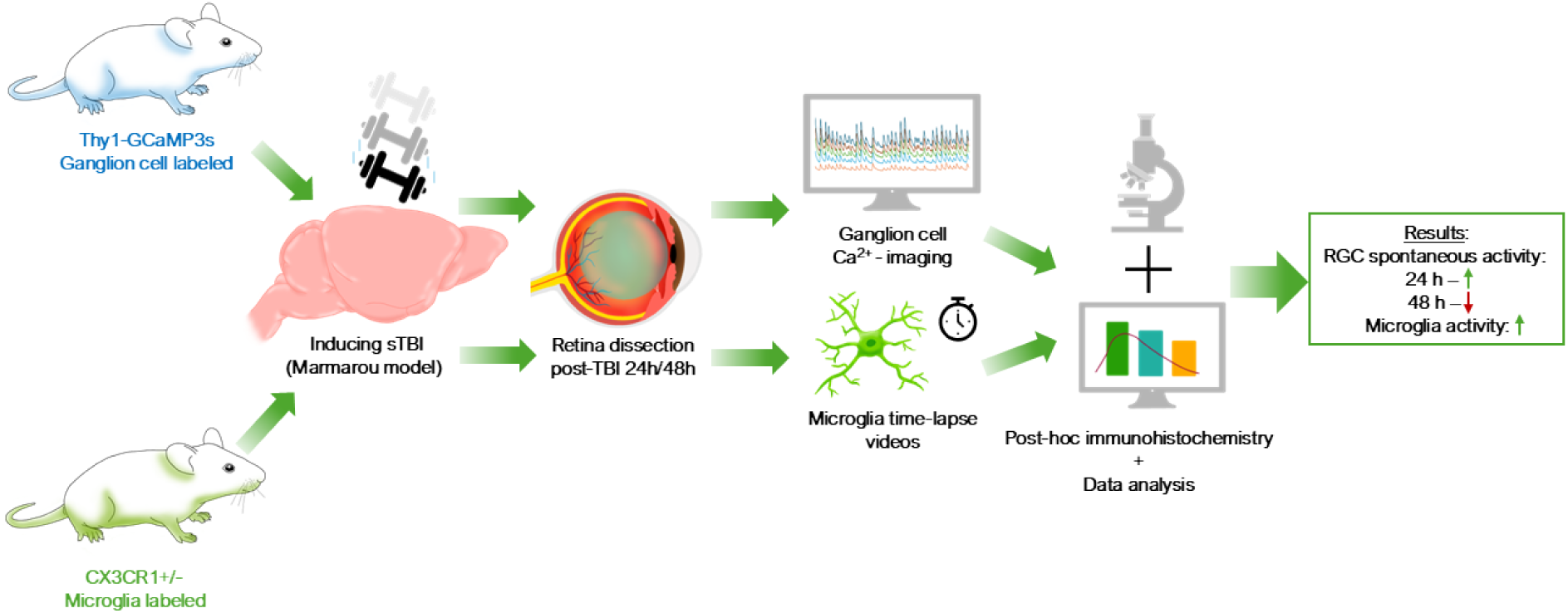

## 1. Background

Traumatic brain injury (TBI) is a serious public health problem (https://doi.org/10.1016/S1474-4422(17)30371-X). It is a collective term for all injuries sustained when the scalp is struck by an external force. Hard falls, traffic-related incidents, and physical assault are a few examples of accidents that could happen to anyone and could result in significant and long-term health issues. Depending on the severity of the trauma, TBI can be categorized as either mild (commonly called as concussion), moderate, or severe. TBI often results in sensory, cognitive, and emotional impairments, and affected individuals may experience difficulty returning to their normal daily activities, including work. Primary symptoms, such as dizziness, nausea and headaches, speech difficulties, hearing ringing in the ears, and skull fractures, typically arise immediately after the injury. However, the impact of TBI extends beyond these initial symptoms, as secondary injury processes may develop hours later, including edema, ischemia, and hypoxia, or even years after the initial trauma https://doi.org/10.1038/nrdp.2016.84).

Beyond its primary effects on the brain, TBI can also affect anatomically connected structures, including the optic nerve and retina. The mechanical forces and pressure waves generated during injury can compress and stretch neural tissue, potentially damaging the optic nerve and visual pathways. It can trigger autonomic dysfunction and an inflammatory response, promote apoptosis, and cell death. Various symptoms accompanying TBI have an effect on the visual system, with examples such as double vision, blurred vision, reduced peripheral vision, pupillary reflex damage, photosensitivity, and optic nerve abnormalities (https://doi.org/10.1515/revneuro-2018-0015).

Microglia, the resident immune cells of the CNS, play a crucial role in responding to TBI and in the context of altered neuronal spontaneous activity. Their dynamic responses to injury are characterized by activation, proliferation, and polarization, which significantly influence the neuroinflammatory environment and subsequent neuronal outcomes. Immediately following TBI, microglia activate and undergo morphological changes, transitioning from a ramified resting state to an amoeboid active state. This activation is a critical early response aimed at managing the acute inflammatory process. Microglia respond by producing cytokines such as tumor necrosis factor-alpha (TNF-α) and interleukin-1β (IL-1β), which promote inflammation and neuroprotection but can also lead to secondary neuronal damage if persistently elevated (https://doi.org/10.3892/mmr.2024.13228; https://doi.org/10.1111/cns.12360).

Acute microglial activation is essential for clearing cellular debris and apoptotic cells through phagocytosis, thereby supporting tissue repair and recovery. However, prolonged or chronic activation, which often occurs following severe injury, can sustain a pro-inflammatory state that exacerbates neuronal loss and contributes to cognitive decline (https://doi.org/10.1016/j.bbi.2021.08.210; https://doi.org/10.1016/j.cell.2020.02.013). Microglia exhibit a polarized response following TBI, differentiating into two main phenotypes: the pro-inflammatory M1 phenotype and the anti-inflammatory, repair-associated M2 phenotype (https://doi.org/10.1089/neu.2017.5540; https://doi.org/10.1186/s12974-020-02041-7). Initially, M1 microglia dominate the response to injury, releasing pro-inflammatory cytokines that can promote further neuronal damage if not properly regulated. The transition to M2 polarization is essential for recovery, as M2 microglia produce anti-inflammatory factors and support tissue repair and neuroprotection (https://doi.org/10.1016/j.celrep.2017.04.047; https://doi.org/10.1089/neu.2017.5540).

Following TBI, neuronal spontaneous activity can increase due to dysregulation of the local microenvironment. Activated microglia can contribute to excitotoxicity through the release of inflammatory mediators and reactive oxygen species (ROS), leading to heightened neuronal excitability and spontaneous firing (https://doi.org/10.1016/j.celrep.2017.04.047; https://doi.org/10.2174/1570159x19666211202123322). Elevated neuronal activity, in turn, can further exacerbate inflammation and promote additional microglial activation, creating a feedback loop that complicates recovery and may contribute to additional neuronal apoptosis (https://doi.org/10.1111/cns.12360; https://doi.org/10.1186/s12974-020-02005-x). Studies using animal models have demonstrated that deficits in cognitive performance correlate with the degree of microglial activation and neuronal hyperactivity following injury, highlighting the detrimental impact of this interaction on neuronal function (https://doi.org/10.1016/j.bbi.2021.08.210; https://doi.org/10.3389/fneur.2021.796704).

Prolonged microglial activation can result in chronic neuroinflammation, which contributes to impairments in synaptic plasticity and cognitive function. This sustained inflammatory response complicates recovery from TBI and can result in neuronal apoptosis, long-term neurodegeneration, and behavioral deficits. Research has shown that effective management of neuroinflammation is essential for promoting recovery and minimizing lasting damage (https://doi.org/10.1111/cns.12360;https://doi.org/10.1111/ejn.13959).

Apoptotic processes play a central role in TBI-induced neurodegeneration, and Caspase-3 (Casp3) is a key molecule involved in programmed cell death. As earlier studies have shown, the activation of Casp3 is associated with the activation of microglia. In cases of retinal injury, apoptotic signaling pathways are initiated, which may lead to increased levels of activated Casp3 (10.3390/ijms24054451) . Therefore, it is reasonable to assume that TBI may also result in elevated Casp3 levels in the retina. However, increased Casp3 activation is not exclusively associated with cell death, as microglial cells have been shown to exhibit elevated levels of activated Casp3 without undergoing apoptosis, suggesting additional regulatory or functional roles for this molecule in microglial activation (https://doi.org/10.1038/nature09788).

While the studies described above have shown that TBI causes neuroinflammation, microglial activation, and RGC degeneration in the retina, these findings largely reflect subacute or chronic time points after injury. In contrast, the very early phase following severe TBI remains poorly described. Specifically, detailed knowledge of how retinal microglia respond in the first 24–48 h, how RGC activity functionally changes during this acute window, and when apoptotic pathways are initiated is lacking. Furthermore, these early events have rarely been examined under controlled ex vivo conditions, leaving a critical gap in our knowledge of the immediate cellular and functional responses of the retina to severe TBI.

Precisely for this reason, the aim of this study was to investigate microglial activation and apoptosis at 24 and 48 hours following TBI in a mouse model compared to a control group. This approach allowed us to examine the temporal progression of injury-induced pathological changes. In addition, we aimed to evaluate the physiological changes occurring in retinal ganglion cells following TBI and to determine whether these changes are associated with increased Casp3 activation. We aimed to assess whether microglial activation occurs after TBI, which would serve as an indicator of ongoing inflammatory processes.We hypothesized that severe TBI induces a rapid retinal innate immune response characterized by early microglial activation, altered microglial motility, transient RGC hyperactivity, and caspase-3 upregulation within the first 48 h post-injury.

## 2. Materials and methods

### 2.1 Ethics statement

All animal procedures were conducted in accordance with the European Union Directive 2010/63/EU on the protection of animals used for scientific purposes and followed the ARRIVE guidelines. Experimental protocols were approved by the Institutional Animal Care and Use Committee of the University of Pécs (approval number: BA02/2000-31/2023). All efforts were made to minimize animal suffering and to reduce the number of animals used.

### 2.2 Animals

Two different mouse strains were used in the experiments. Thy1-GCaMP3 transgenic mice were used to label retinal ganglion cells. Adult (2 months to 11 months, n=7) Thy1-GCamP3 (source: JAX #029860), male mice were maintained in a 12/12 hour dark/light cycle; all experiments were carried out during the day with dissections between 9–11 AM, non-dark-adaptated prior to experiments. To study microglia, we used Cx3Cr1(+/-)GFP mice (4 months to 11 months, n=22), held under similar conditions, in which microglia can be selectively identified based on GFP expression driven by the CX3CR1 receptor promoter. The line was a kind gift from Adam Denes’s group and extensively characterised before (10.1038/jcbfm.2008.6, 10.1038/s41467-024-49773-1); originally obtained from the European Mouse Mutant Archive (EMMA), backcrossed to C57Bl6J (Jung et al, 2000).The use of the two mouse strains allowed for the parallel, targeted study of neuronal and immune cell components of the retina. 4 mice were placed in one cage with food and water ad libitum.

### 2.3 Induction of TBI

The first step of the experiment was the induction of TBI. The mouse models were anesthetized with 5% isoflurane with 70:30 N_2_:O_2_ gas mix. After the anesthesia was stabilized, we fixed the mice heads in a stereotaxic frame and used 2% isoflurane in the same gas mix. A midline incision was made in the skull of the animal, and the periosteum was removed. Then, a helmet plate was glued between the lambda and bregma sutures. After that, the animal was placed on a sponge bed in a prone position. Thus, the helmet was positioned just below the plastic tube. The copper cylinders were set at the right height to induce the type of TBI we wanted to induce. Among the types of TBI, we wanted to examine severe brain injury, which was performed according to the Marmarou model with a free weight drop of 100 g from a height of 75 cm (https://doi.org/10.1007/978-1-60327-185-1_34). After inducing TBI, the helmet was removed from the animals’ heads, and the wounds were sutured to allow them to recover properly. During the surgical procedure, the physiological parameters of the animals were monitored using a pulse oximeter (MouseOx Plus, Starr Life Corp., Oakmont, PA, USA). Body temperature was monitored using a Homothermic Monitoring System (Harvard Apparatus, Holliston, MA, USA) and maintained at the required 37°C for the animals using a heating pad on the same device. According to the TBI protocol, we tested 2 TBI-treated groups in addition to the control group. The dissection was performed after 24 hours in one group and 48 hours in the other, so that the differences due to time since TBI could be better understood. In all groups, randomly one eye of the animals was examined by calcium ion imaging and the other by post-hoc immunohistochemistry. Left and right eye allocation was randomized. Statistical analyses treated the animal, not the eye, as the biological replicate

### 2.4 Functional Calcium-imaging

During the experiment, retinas from both control and TBI-treated animals were examined using Ca^2+^ imaging. Following TBI induction, the retinas of one group of animals were isolated after 24 h and the other group after 48 h. The imaging process was the same for all three groups. The animals were sacrificed after anesthesia with isoflurane. The eyes were then enucleated, and the cornea was cut below the ora serrata so the lens could be removed from the eye cup. The retinas were carefully isolated and placed in a specialized sample holder under the microscope. A physiological solution continuously perfused the samples at the appropriate temperature throughout the experiment, thereby maintaining physiological conditions.

Ringer’s isotonic saline was used during imaging. The solution was freshly prepared before each experiment. Its composition in 500 ml of distilled water was as follows: 3.6525 g NaCl; 0.1180 g KCl; 0.1417 g CaCl_2_; 0.0862 g NaH_2_PO_4_; 0.1017 g MgCl_2_; 0.9008 g glucose; 1.0501 g NaHCO_3_. While continuously stirring 400 ml dH_2_O, the measured substances were added one by one, except for NaHCO_3_. Before that, the solution was oxygenated with carbogen (95% CO_2_, 5% O_2_) for 10 minutes to prevent the components from contacting each other. Finally, NaHCO_3_ was added, and the solution volume was adjusted to 500 ml. The prepared Ringer’s solution was then continuously oxygenated throughout the experiment.

Imaging was performed using a Nikon Eclipse FN1 upright microscope equipped with a Sony Alpha 6400 camera, which enabled proper adjustment of illumination conditions and video recording. Recordings were acquired at a sampling rate of 50 Hz using a 40× objective. During measurements, the retina was illuminated with 470 nm excitation light, also served as visual stimulus, for 4 seconds after the start of the recording, followed by a 2-second dark period. This cycle was repeated three additional times to determine whether light-evoked responses could be detected in retinal ganglion cells. Subsequently, continuous illumination was applied until the end of the recording, which lasted a total of 90 seconds. During illumination, spontaneous cellular activity was monitored, as indicated by transient fluorescence signals. Ca²⁺ events were processed with ImageJ, FIJI (ΔF/F thresholded, field size 200 µm²)

### 2.5 Ex vivo time-lapse videos

Time-lapse imaging was performed on ex vivo-prepared retinas. Following dissection, retinas from Cx3Cr1(+/-)GFP transgenic mice were placed in carbonated Na-HEPES-Ringer solution (composition per 1000 ml of distilled water: 8.01 g NaCl; 0,1864 g KCl; 2,5 ml CaCl_2_; 1 ml MgCl_2_; 5,044 g glucose; 2,603 g Na-HEPES; pH 7,4) and incubated at 33 °C with continuous perfusion throughout the measurement. The retinas were positioned on the stage of a fluorescent microscope (Nikon Eclipse FN1 Upright microscope, equipped with a Sony Alpha 6400 camera) and images were taken with a 40x objective. Time-lapse imaging was conducted over a 20-minute period, with images recorded at 30-second intervals.

### 2.6 Immunohistochemistry

Following either the Ca²⁺ imaging or time-lapse experiments, retinas were fixed in 4% PFA (4% paraformaldehyde in PBS: 137 mM NaCl; 2.7 mM KCl; 10 mM Na_2_HPO_4_ · 7H_2_O, pH 7.4) for 20 minutes. After fixation, retinas were washed three times for 10 minutes each in PBS. To flatten the tissue, the retinas were placed on the surface of the sample holder and blocked in CTA (5% Chemiblocker, 0.5% TritonX-100, 0.05% Na-azide in PBS) overnight. The following day, a solution containing the appropriate proportion of primary antibodies was prepared. The primer antibodies were guinea pig Iba-1 (SySy Antibodies; #234004; 1:1000) and rabbit Casp3 (Novus Biologicals; #AF835; 1:500). CTA was also used for dilution. The solution was vortexed and centrifuged, then pipetted onto retinas, which were incubated for 48 hours at room temperature.

After primary antibody incubation, the retinas were washed three times in PBS for 10 minutes. During this period, the secondary antibody–CTA mixture was prepared. The secondary antibodies were anti-guinea pig Alexa 647 (Invitrogen; #A21450; 1:500) and anti-rabbit Cy3 (Jackson ImmunoResearch; #715-165-150; 1:500). After vortexing and centrifugation, the secondary antibodies were applied to the retinas and incubated at room temperature for 24 hours.

Following secondary antibody incubation, the retinas were washed three more times for 10 minutes each in PBS, mounted on slides, and covered with Vectashield cover slips (Vector Laboratories). Prepared samples were imaged using a Zeiss LSM 710 confocal laser scanning microscope, employing 6x and 20x objectives.

### 2.7 Image analysis

All data was processed using ImageJ/FIJI software (v.1.54p, Java 21.0.7). In the case of Ca^2+^ imaging, the video frames were merged together to create a single image, allowing individual retinal ganglion cells to be identified and marked. These were defined as “ROIs” (Regions of Interest) by the software, which then restricted subsequent analysis to these regions within the video recordings. The ROIs selection was manual. Based on the activity of the ganglion cells, a value was assigned to each frame, with higher numbers of active cells corresponding to higher values. These numbers were exported to Microsoft Excel and plotted as line graphs to illustrate both light-evoked responses and spontaneous activity for each cell. Additionally, the number of frames per minute at a 50 Hz acquisition rate was calculated to enable comparison between control and brain-injured groups. The resulting curves reflected the relative Ca²⁺ activity over the measured time interval.

During the time-lapse recordings, images were acquired for 20 minutes at 30-second intervals. The resulting image sequences were compiled into a time-lapse video in FIJI, and individual TIFF files were generated. Microglial cells were manually selected in the TIFF files using the MotiQ Fiji plugin (10.1091/mbc.E21-11-0585), and the motion analysis was subsequently performed for each cell with the plugin.

The confocal microscope images were adjusted for antibody signal intensity in each channel. In retinas that did not originate from CX3CR1(+/-)GFP mice, Iba-1 staining was applied, and a noise-canceling procedure was used to remove background signals. Microglial cells of the retina are located in superficial (ganglion cell layer - GCL and inner plexiform layer - IPL) and deep layers (outer plexiform layer - OPL), which were analyzed separately based on the Z-axis of the confocal images. Microglia were classified according to morphological features as either activated or non-activated using the MotiQ plugin. Cells were classified as activated when ramification index fell below X. For Casp3 image analysis, samples were similarly divided into superficial and deep layers, which were evaluated independently. Casp3 labeling was identified as discrete, punctate staining. For each layer, the total number of Casp3 staining was quantified, and then the amount of dots that localized to microglia cells was determined separately. To this end, Casp3 labeling was evaluated together with microglia markers, and only those stainings that showed clear colocalization with microglia cells were classified as microglia-associated.

### 2.8 Statistics

Statistical analyses were performed using JASP software (v0.95). Quantitative data obtained from image analyses in Fiji program were first exported to Microsoft Excel format, and then the data files were imported into JASP for statistical evaluation. Data distribution was assessed using the Shapiro–Wilk test for normality, and homogeneity of variances was evaluated using Levene’s test. Although deviations from normality were observed in some data sets, parametric statistical analysis was used because one-way analysis of variance (one-way ANOVA) is generally considered appropriate for data sets that deviate slightly from normal distribution, especially when group sizes are comparable. Accordingly, one-way ANOVA was used to compare the individual data groups. After the detection of a significant main effect, Tukey’s post hoc test was performed to reveal the differences between the individual groups. The unit of analysis differed across experiments: microglial analyses were based on individual cells, ganglion cell recordings were evaluated based on spike counts, and caspase staining was quantified by counting individual dots. During the analyses, the significance level was determined as p < 0.05 (significant, *), p < 0.01 (highly significant, **) and p < 0.001 (very highly significant, ***). Boxplot diagrams were prepared to illustrate the results.

## 3. Results

### 3.1 Spontaneous activation of ganglion cells

We investigated the spontaneous activity of retinal ganglion cells by quantitative analysis of the number of Ca²⁺ events occurring in the cells, as it is known from previous studies that neuronal spontaneous activity may increase during nervous system injuries or inflammatory processes (https://doi.org/10.1186/s12974-021-02236-6, https://doi.org/10.1038/s41467-023-36093-z). Changes in ganglion cell Ca²⁺ activity are therefore be sensitive indicators of the development and time course of pathological conditions.

The results of the analysis showed that 24 hours after traumatic brain injury (n=2), the number of spontaneous Ca²⁺ events in ganglion cells significantly increased compared to the control group (n=3) **(Fig.1)**. This increased activity is presumably part of the nervous system response characteristic of the acute phase following injury, which may be due to the release of inflammatory mediators, changes in cell excitability, and neuron–glia interactions mediated by microglia activation. This transient hyperactivity is consistent with inflammation-driven neuronal hyperexcitability reported in other neuroimmune injury models.

**Figure 1.**
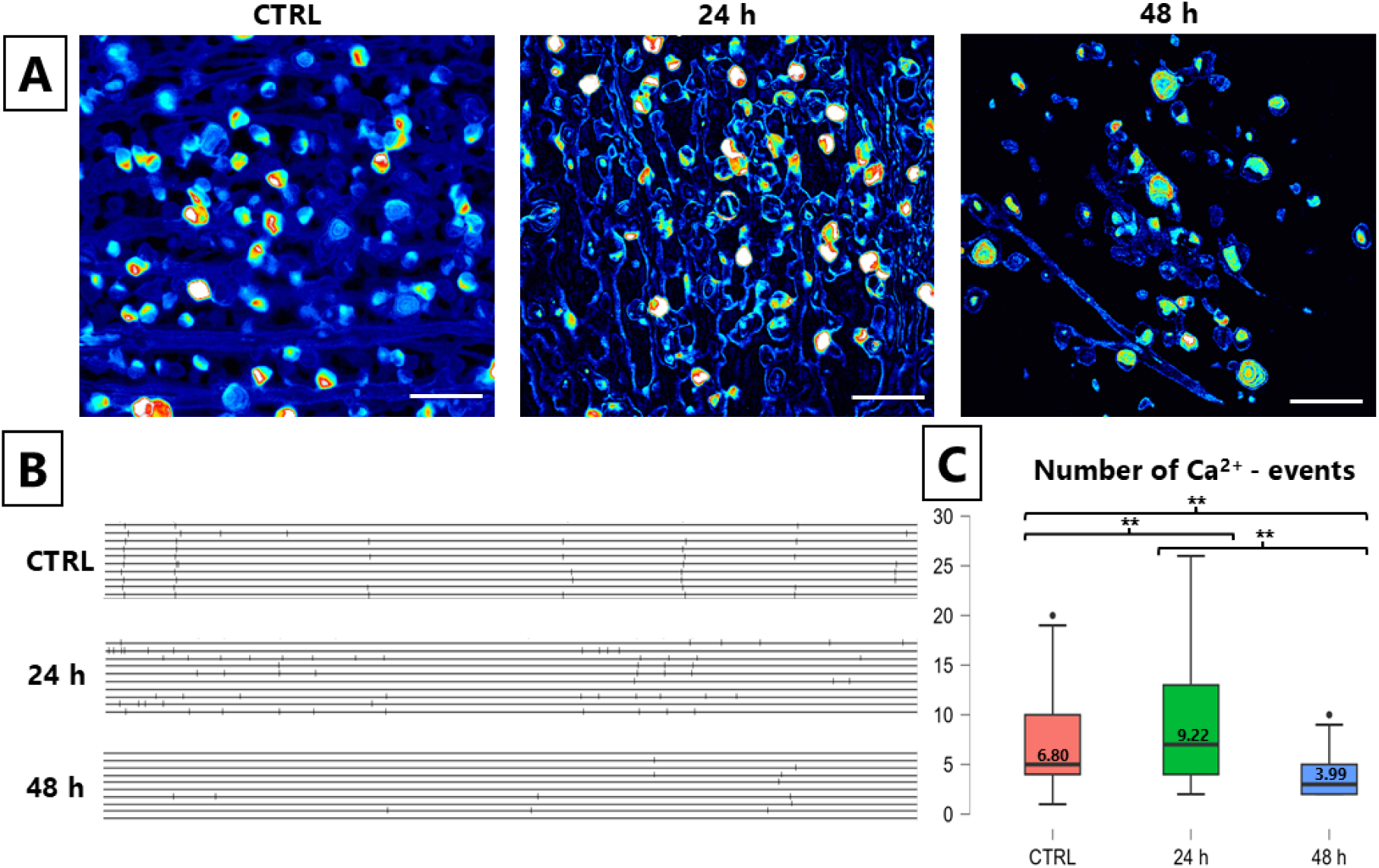
Changes in spontaneous activity following TBI in RGCs. A) Representative calcium imaging images from the CTRL, 24 h and 48 h post-TBI groups. The color scale indicates cell activity, with warmer colors (yellow–red) corresponding to higher activity. B) Representative plots showing calcium events from 10–10 cells per group. Vertical lines indicate individual calcium transients. C) Quantitative analysis of the averaged calcium event frequency across all retinas examined in boxplot form. Both the representative images and the averaged data show that cell activity significantly increases at 24 h post-TBI compared to control and decreases at 48 h. This suggests that retinal activity increases transiently and then declines later. (n=7; total cell number=305). Tukey’s post hoc ** p<0.01

In contrast, 48 hours after TBI (n=2), the number of Ca²⁺ events decreased significantly compared to the 24-hour time point. This decrease suggests that the functional state of ganglion cells deteriorates after the initial hyperactive state. We hypothesize that this phenomenon may be related to cell damage and cell death occurring after injury, which results in a decrease in the number of ganglion cells. As a result, the measured spontaneous Ca²⁺ activity becomes lower.

Overall, our results suggest that the spontaneous Ca²⁺ activity of ganglion cells changes dynamically in the period following TBI. The increased activity is observed in 24-hrs, while later the decline of neuronal functions dominates. These results support the assumption that the examination of ganglion cell Ca²⁺ activity may be a suitable method to monitor pathological processes after injury and may contribute to a better understanding of the time course of neuronal damage.

### 3.2 Activation of microglia

Following the demonstration of increased apoptosis and altered spontaneous activity of ganglion cells, the next step was to investigate the immune cells of the retina, microglia, as they play a key role in the development of the inflammatory response and cellular damage following injury. Microglia are present in all layers of the retina, but they typically occur in two distinct layers based on their distribution. Therefore, in our studies, we focused on microglia populations located in the superficial (GCL+ IPL) and deeper (OPL) layers of the retina.

The degree of microglia activation was characterized by determining the ramification index of the cells, which reflects the morphological differences between the resting and activated states. Resting microglia cells are characterized by small cell bodies and finely branched processes, while activated microglia cells have enlarged cell bodies and reduced, amoeboid-like process structures, consistent with previous observations by Davis et al (https://doi.org/10.1038/s41598-017-01747-8).

In control retinas, microglia were predominantly ramified and evenly distributed across IPL and OPL layers. Analysis of the superficial layer showed significant changes after TBI **(Fig.2)**. The ramification index decreased significantly 24 hours after TBI (n=5), indicating activation of microglia. At 48 hours after injury (n=4), the ramification index partially recovered, but the values still differed from the control group (n=4), suggesting that microglia activation still existed at this time point, albeit in a more moderate form.

**Figure 2.**
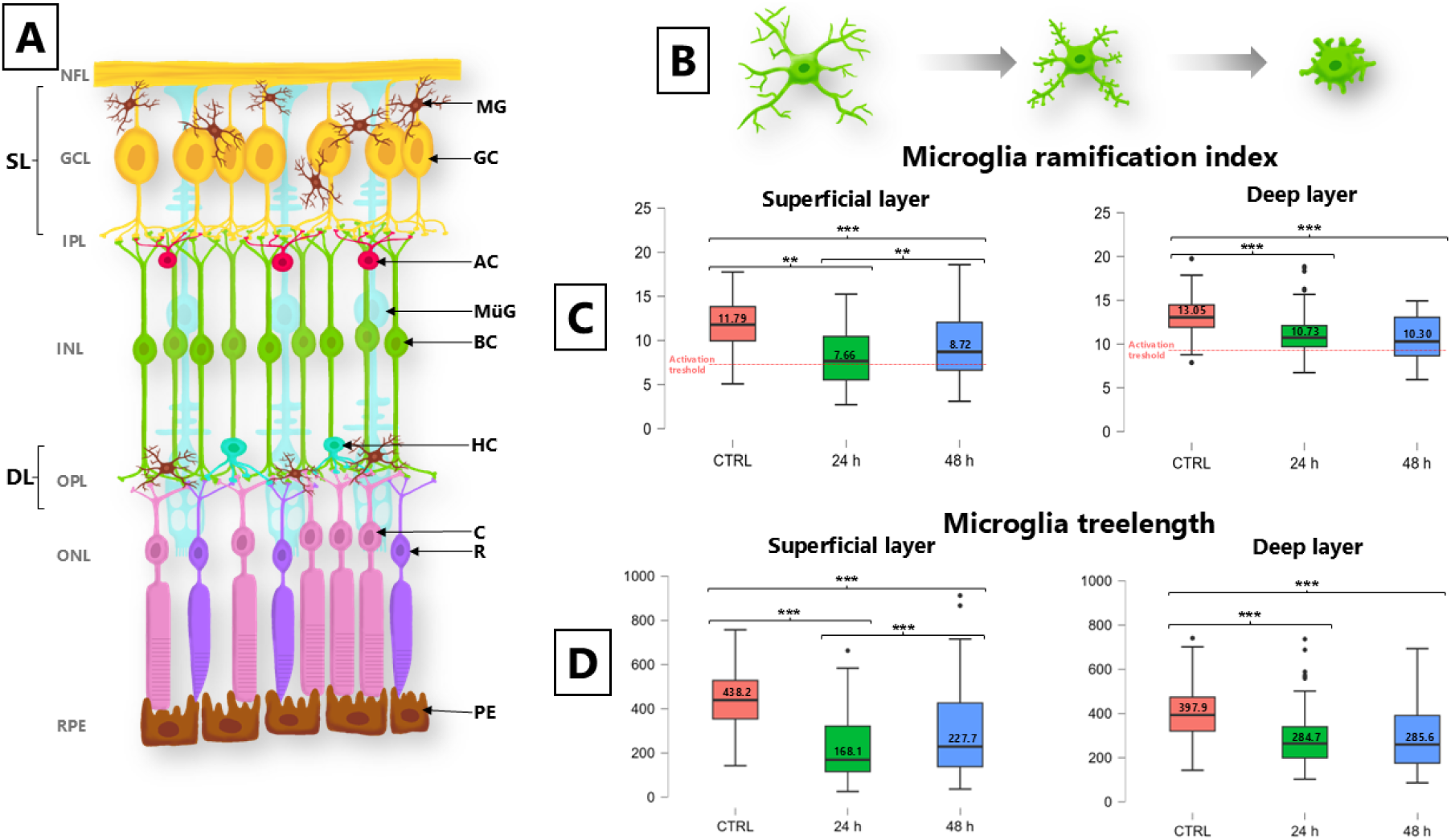
Morphological changes in retinal microglia after TBI. (A) Schematic representation of the mouse retina, showing the main layers and cells. (B) Representation of microglia activation. In the resting state, microglia cells have long, highly branched processes, while during activation, the processes gradually shorten, retract, and the morphology of the cells becomes more compact. (C) Quantitative analysis of the microglia ramification index in the superficial and deep retinal layers. The red dashed line indicates the activation threshold below which cells are considered activated. Since activation is associated with the retraction of the processes, a lower ramification index indicates increased microglia activation. (D) Quantitative analysis of the microglia tree length, a measure of the branching and morphological complexity of the cells. TBI results in a significant decrease in microglia ramification index and total process length in both the superficial and deep retinal layers, indicating activation of the cells. The most pronounced changes are observed 24 hours after injury, while 48 hours later there is a partial recovery, but the values are still shifted towards the activated state compared to the control group. Tukey’s post hoc **p<0.01; ***p<0,001 (n=13; total cell number=333) SL: superficial layer, DL: deep layer, NFL: nerve fiber layer, GCL: ganglion cell layer, IPL: inner plexiform layer, INL: inner nuclear layer, OPL: outer plexiform layer, ONL: outer nuclear layer, RPE: retinal pigment epithelium, MG: microglia, GC: ganglion cell, AC: amacrine cell, MüG: Müller glia, BC: bipolar cell, HC: horizontal cell, C: cone, R: rod, PE: pigment epithelial cell

A similar trend was observed in the deeper layer, but the activation pattern showed different dynamics. Here, too, significant changes in the ramification index occurred after TBI, indicating activation of microglia. The differences between the superficial and deep layers suggest that the microglia response is layer-specific, which is consistent with previous studies in rats that described differences in the ratio of activated to total microglia populations in different layers of the retina (https://doi.org/10.3390/ijms24054451).

### 3.3 Caspase activation

To map the cellular damage processes following TBI, we examined the extent of apoptosis in the retinas by immunohistochemical labeling of activated caspase-3 (simply referred as Casp3). Casp3 is a key enzyme in apoptosis, therefore the appearance of its activated form is a reliable indicator of the process of programmed cell death. During the analyses, Casp3-positive staining was quantified both in all cell populations and exclusively in microglia cells, taking advantage of microglia-specific labeling. The studies were performed separately in the superficial and deep layers of the retina.

In the superficial layer, the number of Casp3-positive cells, considering all cells, increased significantly 24 hours after TBI (n=6), and then increased even further 48 hours after injury (n=6) compared to the control group (n=5) (Mean values: SL: CTRL - 1; post-TBI 24 h - 11; post-TBI 48 h - 28; DL: CTRL - 9.5; post-TBI 24 h - 15; post-TBI 48 h - 36, see **Fig.3**). A similar trend was observed in the analysis performed only in microglia: the number of Casp3-positive microglia cells increased as early as 24 hours after TBI, which continued to increase at 48 hours (Mean values: SL: CTRL - 0; post-TBI 24 h - 4; post-TBI 48 h - 18; DL: CTRL - 7.5; post-TBI 24 h - 9; post-TBI 48 h - 31). This suggests that in the superficial layer, not only neuronal and other cell types are affected by apoptosis after TBI, but also a portion of microglia cells has an increased expression pattern of Casp3.

**Figure 3.**
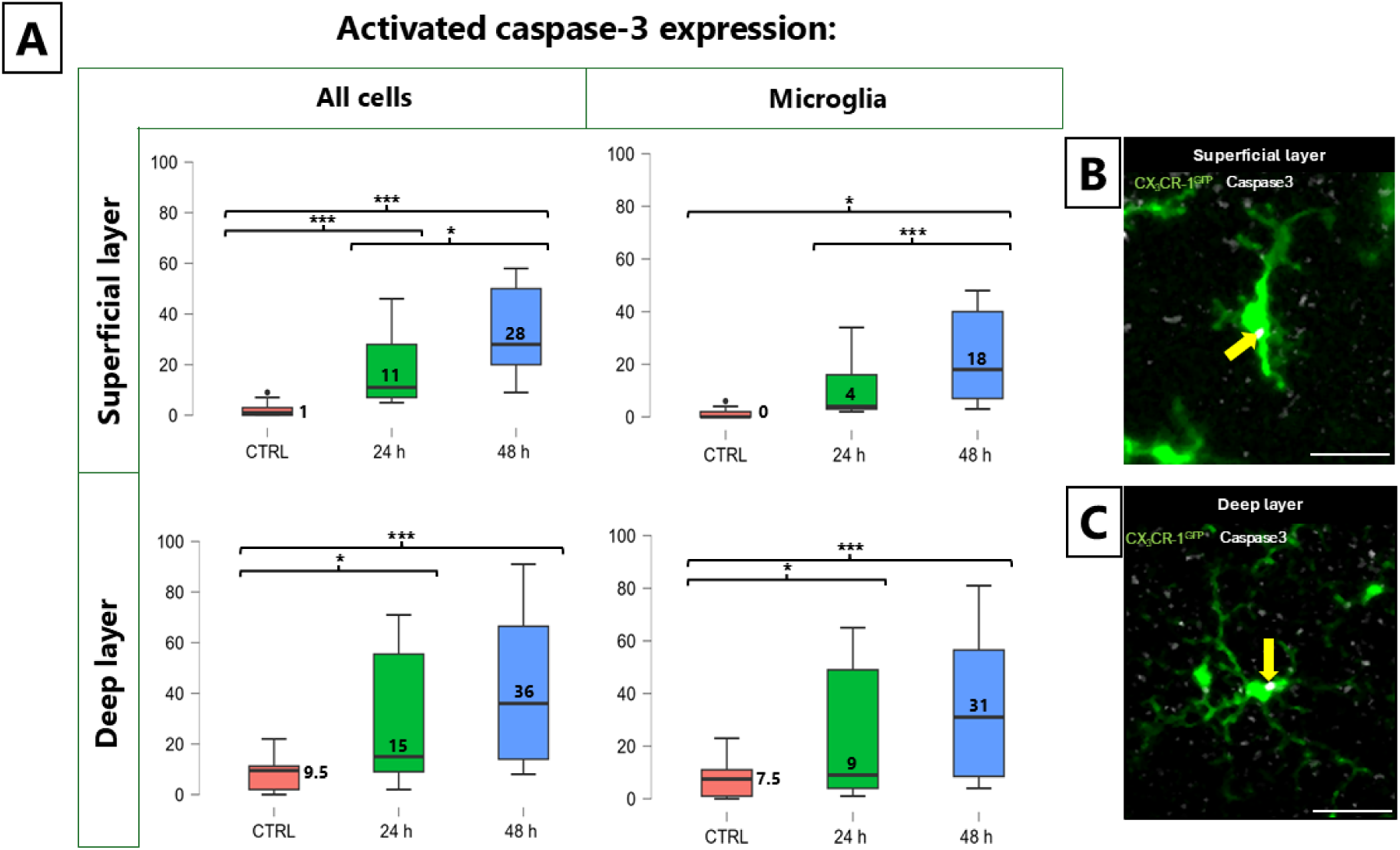
Traumatic brain injury-induced caspase-3 activation in retinal cells and microglial population. (A) Quantitative analysis of the number of cleaved caspase-3 positive cells in the superficial and deep layers of the retina. The figures show separately the total caspase-3 positive cells and the caspase-3 expression associated with microglia in the CTRL, 24-hour and 48-hour groups. (B) Representative confocal image of a microglia from the superficial layer 24 hours after traumatic brain injury. (C) Representative confocal image of a microglia from the deep layer 24 hours after traumatic brain injury. After TBI, the number of activated caspase-3 positive cells significantly increased in both the superficial and deep retinal layers compared to the control. The increase was observed not only in the total cell population but also in microglial cells, and the values were increased beyond 48 hours compared to the 24-hour group. Representative images confirm the presence of activated caspase-3 in microglial cells. The results indicate progressive apoptotic processes in both layers of the retina following traumatic brain injury. Yellow arrows: activated-caspase3 marker. Scale: 25 μm. Tukey’s post hoc *p<0.05; ***p<0.001 (n=17)

### 3.4 Temporal changes in microglial motility following TBI

Changes in microglia motility were examined by analyzing time-lapse recordings after TBI. During the analyses, we focused exclusively on the superficial microglial layer of the retina, while the deeper layer was not included in the study. The reason for this was that in the later planned in vivo studies, only the superficial microglial layer can be visualized for technical reasons, so the present results provide a direct basis for comparison for future in vivo measurements.

Analysis of the ramification index did not show significant differences compared to the control group either 24 or 48 hours after TBI, indicating that the global morphological complexity of microglial cells did not change significantly within this time window. However, clear differences were observed in the dynamics of microglial processes based on the time-lapse recordings, which prompted a more detailed quantitative analysis of motility . The time lapse images showed a pronounced movement of MG endfeet **(Fig.4A, Supplementary Video 1)**. Time-color-coded images from time-lapse videos show that the processes extend and retract repeatedly, and also maintain a continuous scanning activity (**Fig.4B).**

**Figure 4.**
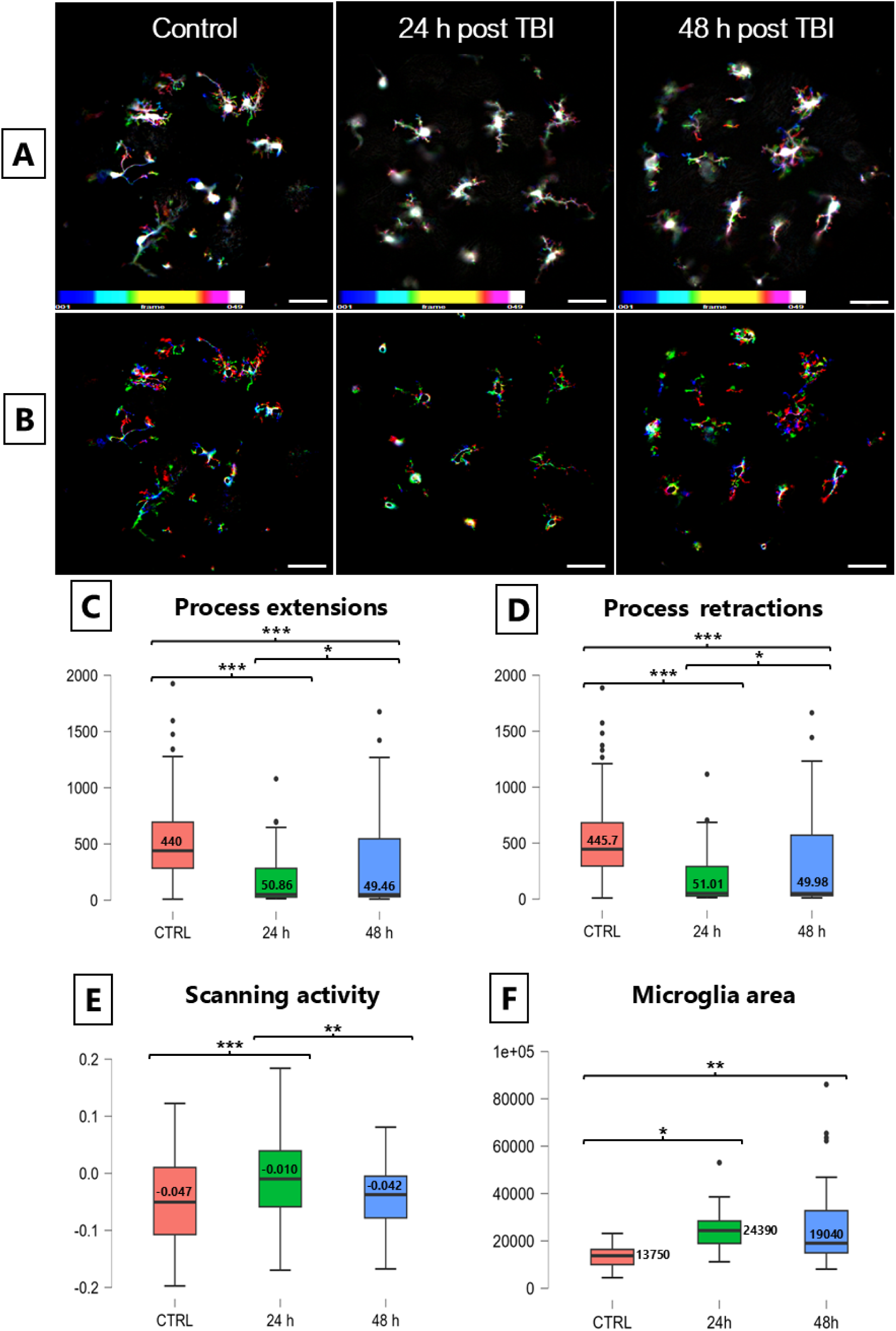
Time-lapse microscopy and microglia motility parameters after traumatic brain injury. (A) Representative time-lapse images illustrating changes in microglial motility after injury. The images were generated by collapsing 51 frames from a 20-minute time-lapse recording (25 s) for each group. The colors correspond to individual frames (see color scale in the lower left corner). Rainbow-colored structures indicate time-dependent positional changes, whereas static structures accumulate all colors and therefore appear white. These images demonstrate that following TBI, microglia show increased motility. (B) Differential images of the dynamic changes in the processes and cell area in the same samples, where different colors separate the temporal movement of the cell boundaries and the phases of the extending and retracting areas. (C) Average extensions of microglial processes based on quantitative analysis of time-lapse recordings. A significant decrease is observed after TBI compared to the control group. Y-axis: extension of processes per minute. (D) Average retractions of microglial processes. A significant decrease is observed at 24 hours, while partial normalization is observed at 48 hours after injury. Y-axis: processes withdraw per minute. (n=18; total cell number=270) (E) Changes in microglia dynamic fraction of projected area. Scanning activity characterizes the dynamic change in the area covered by cells over time (20 min.). At 24 hours after TBI, a significant increase is observed compared to the control, while by 48 hours the values show a decrease, approaching the control level. (n=18; total cell number=270) (F) Changes in microglial cell area. The average cell area increased significantly 24 hours after TBI compared to control, indicating microglial activation. At 48 hours after injury, the values decreased compared to 24 hours, but remained elevated compared to control. Y axis: cell area in pixel units (pixels²) Scale: 50 μm. (n=14; total cell number=70) Tukey’s post hoc *p<0.05; **p<0.01; ***p<0.001

To characterize overall dynamic activity change in TBI, we first analyzed the dynamic fraction of the projected area (scanning activity in MotiQ) **(Fig.4 E)**. This parameter reflects the extent of endfeet-related movements over time (20 min.). At 24 hours after TBI (n=6), a significant increase was observed compared to the control group (n=7), while at 48 hours (n=5)the values decreased, approaching control levels. To further dissect microglial motility, we quantified the extension and retraction of cellular processes **(Fig.4 C and D)**. The results showed that 24 hours after TBI (n=6), both the mean extension and retraction rates were significantly reduced compared to the control group (n=7). This reduction indicates an activated microglial state characterized by shorter and less dynamic processes. At 48 hours after TBI (n=5), partial normalization of these parameters was observed: although motility remained altered compared to control, both extension and retraction rates showed recovery relative to the 24-hour time point.

In addition to dynamic parameters, we also examined changes in cell morphology by measuring the average cell area **(Fig.4 F)**. Cell area was measured in FIJI using manually defined ROIs and is reported in pixel units (pixels²). A significant increase in cell area was detected 24 hours after TBI (n=4), indicating microglial activation. At 48 hours after injury (n=6), the values decreased compared to 24 hours but remained elevated relative to the control group (n=4). Taken together, these results demonstrate that microglial activation after TBI is a dynamic and time-dependent process. At 24 hours post-injury, microglia exhibit increased cell area and reduced process dynamics, reflecting a pronounced activated state. By 48 hours, partial normalization of these parameters occurs; however, microglial motility and morphology have not yet fully returned to baseline levels.

## 4. Discussion

### 4.1 Early transient hyperexcitability of retinal ganglion cells after TBI

Spontaneous activity of retinal ganglion cells (RGCs) is essential for retinal circuit organization and functional stability. While prominently studied in development as retinal waves driven by cholinergic amacrine cells (https://doi.org/10.1371/journal.pone.0077658; https://doi.org/10.1523/jneurosci.23-19-07343.2003), aberrant spontaneous RGC activity also emerges in adult pathological states. Increased RGC firing has been reported in glaucoma and optic nerve injury, where it is associated with maladaptive hyperexcitability and subsequent neurodegeneration (Weitlauf et al., 2014; Cheng et al., 2021; Risner et al., 2018).

In this study, severe TBI induced a pronounced increase in spontaneous RGC Ca²⁺ activity at 24 hours, followed by a decline below control levels at 48 hours. This biphasic response suggests an acute phase of injury-induced hyperexcitability that rapidly transitions toward functional impairment. Neuroinflammatory signaling likely contributes to this shift, as activated glial cells release cytokines such as TNF-α that enhance RGC excitability via upregulation of voltage-gated sodium channels, including NaV1.6 (Cheng et al., 2021). Early inflammatory remodeling can promote aberrant oscillatory firing (Hüll et al., 2019; Li et al., 2022), whereas sustained inflammation and impaired trophic support are associated with declining spontaneous activity and delayed RGC death (Berkelaar et al., 1994; Villegas-Pérez et al., 1993; Kim et al., 2013). Together, these data indicate that early retinal hyperexcitability after TBI is a transient state preceding neurodegenerative dysfunction.

### 4.2 Rapid microglial activation across retinal layers following TBI

Microglia play a central role in maintaining synaptic integrity and retinal homeostasis, and their dysfunction contributes to synaptic disorganization and visual impairment (Wang et al., 2016). We observed robust microglial activation in both superficial and deep retinal layers within 24 hours after severe TBI, with activation persisting at 48 hours. This widespread response indicates that TBI elicits a global retinal inflammatory reaction rather than a layer-restricted effect. Layer-specific differences in activation dynamics likely reflect distinct synaptic environments and injury sensitivity within the retinal circuitry.

### 4.3 Microglial motility reveals sustained functional activation

Beyond static morphological changes, time-lapse imaging demonstrated marked alterations in microglial motility after TBI, particularly during the early post-injury phase. Notably, partial normalization of ramification indices at 48 hours contrasted with persistent abnormalities in process dynamics, indicating that morphological measures alone do not fully capture microglial activation state. Dynamic motility parameters therefore provide critical complementary information on microglial engagement during acute retinal inflammation.

Modulating microglial activation and polarization toward reparative phenotypes has emerged as a promising therapeutic strategy after TBI (Willis et al., 2020; Yang et al., 2024). Pharmacological approaches, including glucocorticoids, can influence microglial responses, although their efficacy and optimal timing remain unresolved (Eyolfson et al., 2020; Lee & Choi, 2018).

### 4.4 Sustained retinal caspase**-**3 activation links apoptosis and neuroinflammation

Caspase-3 is a key mediator of apoptotic signaling and secondary injury following traumatic brain injury. Here, severe TBI induced a robust increase in activated caspase-3 in the retina at 24 hours, affecting both microglia and other retinal cell populations, with elevated levels persisting at 48 hours. Although mean values tended to increase over time, the absence of consistent differences between post-injury time points suggests rapid initiation and maintenance of apoptotic signaling rather than progressive escalation within the first 48 hours.

These findings are consistent with previous reports of early caspase-3 upregulation after TBI (https://doi.org/10.1523/JNEUROSCI.21-19-07439.2001) and chronic persistence of cleaved caspase-3 in injured brain regions (Glushakova et al., 2018). Importantly, caspase-3 activation within microglia may reflect non-apoptotic roles in inflammatory regulation, as microglial inhibition reduces both cytokine production and caspase-3 activity in retinal disease models (Krady et al., 2005). Caspase-3–mediated apoptosis is a common feature across CNS and retinal injury paradigms, including optic nerve axotomy and retinal degeneration (https://doi.org/10.1089/neu.1999.16.153; Crouzier et al., 2021).

## Summary

Severe TBI triggers rapid, coordinated retinal responses characterized by **early RGC hyperexcitability**, **widespread microglial activation with sustained motility changes**, and **persistent caspase**-**3–associated apoptotic signaling**. These findings establish the retina as a sensitive and functionally informative readout of early CNS pathology following traumatic brain injury.

The studies above robustly support the assertion that caspase-3 activation is significantly elevated within microglia and other retinal cells at 24 and 48 hours post-traumatic brain injury, consistent with ongoing apoptotic mechanisms. These findings highlight the vital role of caspase-3 in the apoptotic processes induced by traumatic brain injuries, establishing a basis for potential therapeutic targets aimed at modulating this pathway to promote neuronal survival following TBI.

## 5. Acknowledgement

We thank Zsuzsanna Helyes and Adam Denes for the Cx3Cr1 mouse line. We acknowledge the administrative help provided by the Szentágothai Research Centre. The research was performed in collaboration with the Imaging Core Facility at the Szentágothai Research Centre of the University of Pécs, and supported by the Nano-Bio-Imaging Core Facility. We thank the staff of the Animal Core Facility of the SZRC, University of Pécs.

## 6. Abbreviations

AC: amacrine cell
BC: bipolar
C: cone
Casp3: caspase-3
CNS: central nervous system
DL: deep layer
GC: ganglion cell
GCL: ganglion cell layer
HC: horizontal cell
IL-1β: interleukin-1β
INL: inner nuclear layer
IOP: intraocular pressure
IPL: inner plexiform layer
MG: microglia
MüG: Müller glia
NF-κB: nuclear factor-kappa B
NFL: nerve fiber layer
ONL: outer nuclear layer
OPL: outer plexiform layer
PE: pigment epithelial cell
R: rod
RGC: retinal ganglion cell
RPE: retinal pigment epithelium
SL: superficial layer
sTBI: severe traumatic brain injury
TBI: traumatic brain injury
TNF-α: tumor necrosis factor-alpha

## Notes

### Competing Interest Statement

The authors have declared no competing interest.

